# ATPase and protease domain movements in the bacterial AAA+ protease FtsH are driven by thermal fluctuations

**DOI:** 10.1101/323055

**Authors:** Martine Ruer, Georg Krainer, Philip Gröger, Michael Schlierf

**Affiliations:** B CUBE – Center for Molecular Bioengineering, Technische Universität Dresden, Arnoldstr. 18, 01307 Dresden, Germany; Max Planck Institute of Molecular Cell Biology and Genetics (MPI-CBG), Pfotenhauer Str. 108, 01307 Dresden, Germany; Molecular Biophysics, Technische Universität Kaiserslautern (TUK), Erwin-Schrödinger-Str. 13, 67663 Kaiserslautern, Germany

**Keywords:** ATP-dependent proteases, conformational dynamics, vesicle encapsulation, protein degradation, single-molecule FRET

## Abstract

AAA+ proteases are essential players in cellular pathways of protein degradation. Elucidating their conformational behavior is key for understanding their reaction mechanism and, importantly, for elaborating our understanding of mutation-induced protease deficiencies. Here, we study the structural dynamics of the *Thermotoga maritima* AAA+ hexameric ring metalloprotease FtsH (*Tm*FtsH). Using a single-molecule Förster resonance energy transfer approach to monitor ATPase and protease inter-domain conformational changes in real time, we show that *Tm*FtsH—even in the absence of nucleotide—is a highly dynamic protease undergoing conformational transitions between five states on the second timescale in a sequential manner. Addition of ATP does not influence the number of states nor change the timescale of domain motions, but affects the state occupancy distribution leading to an inter-domain compaction. These findings suggest that thermal energy, but not chemical energy, provides the major driving force for conformational switching, while ATP, through a state reequilibration, introduces directionality into this process. The *Tm*FtsH A359V mutation, a homolog of the human pathogenic A510V mutation of paraplegin (SPG7) causing hereditary spastic paraplegia (HSP), does not affect the dynamic behavior of the protease but impairs the ATP-coupled domain compaction and, thus, may account for protease malfunctioning and pathogenesis in HSP.

## Introduction

Cellular organisms maintain a stable and functional proteome by fine-tuned homeostasis mechanisms that regulate the expression, folding, and degradation of proteins [1]. Key players in the cellular pathways of protein degradation are AAA+ (ATPases associated with diverse cellular activities) proteases [2,3]. These energy-dependent molecular machines remove dysfunctional, misfolded, aggregated as well as no longer needed proteins from the proteome by their specific unfoldase and peptidase activities [4–9]. The importance of this cellular clearance system is eminently reflected in its impairments, as alterations in AAA+-based proteolysis are associated with various dysfunctions in bacteria [10,11] as well as a broad range of neurodegenerative, metabolic, and cancerous diseases in humans [12–16], most frequently caused only by single point mutations in the protease sequence.

One of the prototypic and highly conserved AAA+ proteases in eubacteria, mitochondria, and chloroplasts is the membrane-embedded metalloprotease FtsH [10,11,17]. FtsH forms ring-like hexameric assemblies of monomer subunits exposing a central pore, through which the unfolded protein substrate is translocated. Each monomeric subunit consists of a protease and ATPase domain, which are connected via a hinge region [10]. Recent crystallographic structures of *Thermotoga maritima* FtsH (*Tm*FtsH) without nucleotide and in the presence of ADP revealed a large conformational change between the ATPase and the protease domain upon ADP binding, thus suggesting an ATP-coupled chemo-mechanical cycle that involves a coordinated opening and closing of the two domains between two states [18,19] (Fig. 1a). Even though these structures showcase the large conformational transitions *Tm*FtsH domains can undergo, mechanistic models inferred from the crystal structures often report only on a limited number of conformational states and do not provide dynamic information. Elucidating the underlying reaction mechanism of *Tm*FtsH and other proteases, however, requires kinetic insights for understanding the interplay of ATP binding to conformational changes and, importantly, for elaborating our understanding in protease deficiency-related diseases. For example, the pathogenic mutation A510V of the human *Tm*FtsH-structural homolog paraplegin (SPG7) can cause hereditary spastic paraplegia (HSP) [13,20–22] and is located right at the hinge between the ATPase and protease domain [23], indicating that a relative movement might be impaired. However, our understanding of the dynamic influence of potentially disruptive mutations on conformational changes and ATP coupling remains elusive.

**Figure 1.**
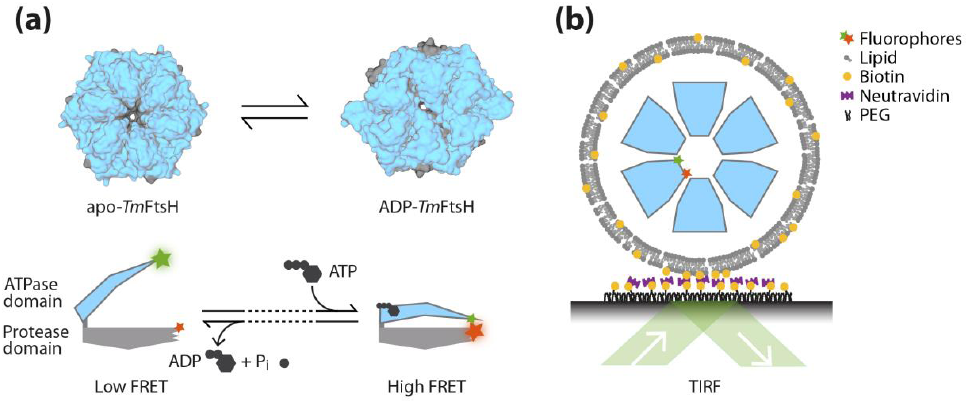
ATPase and protease inter-domain movements in *Tm*FtsH probed by smFRET. (a) Schematics of large-scale conformational changes in *Tm*FtsH. Upper panel: surface representation of hexameric *Tm*FtsH in its apo- (left, pdb: 3kds) and ADP-bound state (right, pdb: 2cea) [18,19]. The protease and ATPase domains are colored in gray and blue, respectively. Visualization was performed in VMD [33]. Lower panel: Schematic of monomer subunit domain conformational changes between an open and a closed state upon ATP binding and hydrolysis as suggested from crystal-structure analysis of *Tm*FtsH. The positions used for labeling with donor (Cy3, green star) and acceptor fluorophores (Cy5, red star) in *Tm*FtsH_184,513_ are indicated. (b) Experimental design of the smFRET assay. Double-labeled *Tm*FtsH_184,513_ monomers were self-assembled in presence of unlabeled *Tm*FtsH to hexamers in DMPC lipid vesicles containing a small fraction of biotin-DPPE. Lipid vesicles were surface-immobilized via a biotin–streptavidin–biotin sandwich and *Tm*FtsH domain conformational changes were monitored by smFRET TIRF microscopy.

Intramolecular dynamics of multi-component enzymes like AAA+ proteases are challenging to resolve because of the experimental difficulties encountered in probing the unsynchronized motions of their constituent subunits or domains [24]. Recent advances in single-molecule experiments have enabled the direct observation of individual molecular machines at work, providing real-time kinetic information on unsynchronized nanoscale motions of enzymes and their subunit structures, which have previously eluded a quantitative description by classical biochemical and structural biology methods [25–29].

Here, we exploited a single-molecule Förster resonance energy transfer (smFRET) approach to study the real-time ATPase and protease inter-domain conformational changes of *Tm*FtsH. We found that inter-domain movements of monomer subunits within assembled *Tm*FtsH hexamers occur on the second timescale, are thermally driven, and weakly coupled to ATP binding or hydrolysis. By performing kinetic analysis based on Hidden Markov modelling, we uncovered five conformational states of *Tm*FtsH, thereby expanding the previous knowledge of the two crystallographic structures. Using this approach, we further studied the effects of the A359V mutation, which is homologous to the human A510V paraplegin mutation, and found that this mutation perturbs the conformational behavior of *Tm*FtsH upon ATP binding, thereby providing a potential mechanism for protease malfunctioning and disease pathogenesis in HSP.

## Results

### Probing the real-time conformational changes in *Tm*FtsH with smFRET

To monitor ATPase and protease inter-domain dynamics in *Tm*FtsH, we established a smFRET assay that allows probing of structural changes in single *Tm*FtsH monomers within self-hexamerized *Tm*FtsH rings. To this end, we created a Cys-light variant (*Tm*FtsH_184,513_) lacking the transmembrane domain of *Tm*FtsH (as described in Bieniossek et al. [18,19]) but carrying two unique Cys residues, one at position 184 in the ATPase domain, and another wild-type cysteine at position 513 in the protease domain. One fraction of *Tm*FtsH_184,513_ was kept unlabeled while the other fraction was reacted with maleimide-functionalized FRET donor (Cy3) and an acceptor (Cy5) fluorophores to obtain double-labeled *Tm*FtsH_184,513_ monomer units (Fig. 1a). The dye labels were placed at positions close to the tip of each domain at a distance such that significant changes in the FRET efficiency are expected if these domains move relatively to each other. Functional assays testing the ATPase and protease activities confirmed that the *Tm*FtsH_184,513_ variant retained both activities (Figs. S1 and S2, respectively). To assemble *Tm*FtsH_184,513_ into their active homohexameric rings, a concentration exceeding the oligomer dissociation constant on the order of ∼400 nM is required. However, working with such high concentrations of labeled protein in solution would result in a high background signal in smFRET experiments [26]. We therefore exploited a lipid vesicle-based nanocontainer approach [30] to increase the effective concentrations of *Tm*FtsH monomers through molecular confinement. We encapsulated a concentrated *Tm*FtsH monomer solution in ∼200-nm diameter phospholipid vesicles, yielding an apparent concentration of the protease monomers inside the vesicles of ∼2.3 mM, thus ensuring self-hexamerization (Fig. 1b). Co-encapsulation of labeled and unlabeled *Tm*FtsH_184,513_ monomers in a 1:5 ratio yields one labeled *Tm*FtsH_184,513_ per assembled hexamer on average, thus allowing to probe conformational switching of one monomer within an active homohexameric ring. The lipid vesicles were composed of 1,2-dimyristoyl-*sn*-glycero-3-phosphocholine (DMPC), which selectively permeabilizes the membrane for ATP addition [30], and contained also a small fraction of 1,2-dipalmitoyl-*sn*-glycero-3-phosphoethanolamine-N-(cap biotinyl) (biotin-DPPE) to tether the vesicles to a quartz slide via biotin–streptavidin interactions. We immobilized the preassembled and *Tm*FtsH_184,513_-filled vesicles on a polyethylene glycol (PEG)–biotin-coated chamber via a biotin–neutravidin–biotin sandwich and performed real-time imaging of fluorescently-labeled single molecules with a prism-based total internal reflection fluorescence (TIRF) microscope [31,32].

### Apo-*Tm*FtsH undergoes sequential transitions between five conformational states

In a first set of experiments, we investigated conformational changes of *Tm*FtsH_184,513_ in the absence of ATP. Donor (*I*_D_) and acceptor (*I*_A_) fluorescence intensities from assembled hexamers in lipid vesicles were recorded for tens of seconds at ten frames per second (Fig. 2a). smFRET time trajectories were extracted by calculating the apparent FRET efficiencies (*E*_app_ = *I*_A_ / (*I*_D_ + *I*_A_)) for each collected data point. Only trajectories exhibiting single bleaching steps of both the donor and acceptor fluorophore were investigated to ensure that FRET efficiency changes arise only from single donor–acceptor labeled monomers and not from multiple co-encapsulated labeled subunits. The donor and acceptor fluorescence time traces showed anti-correlated signals fluctuations between low and high intensity (Fig. 2a), which translate into highly dynamic FRET efficiency changes (Fig. 2b) revealing that *Tm*FtsH_184,513_ exhibits inter-domain movements between the ATPase and protease domain on the second timescale. Based on the structural data, we expected a constant low FRET efficiency in absence of ATP, however, the large distinct changes between *E*_min_ ≈ 0.2 and *E*_max_ ≈ 0.9 observed in the smFRET time trajectories indicate a large hinge-bending motion in the range of ∼3–4 nm between the two domain tips. FRET efficiency histograms from individual time traces (Fig. 2b, right panel) as well as a cumulative FRET efficiency histogram from all trajectories (*n* = 108) resulted in a broad distribution, further supporting that the ATPase and protease domains of *Tm*FtsH_184,513_ dynamically interconvert between multiple states in absence of ATP.

**Figure 2.**
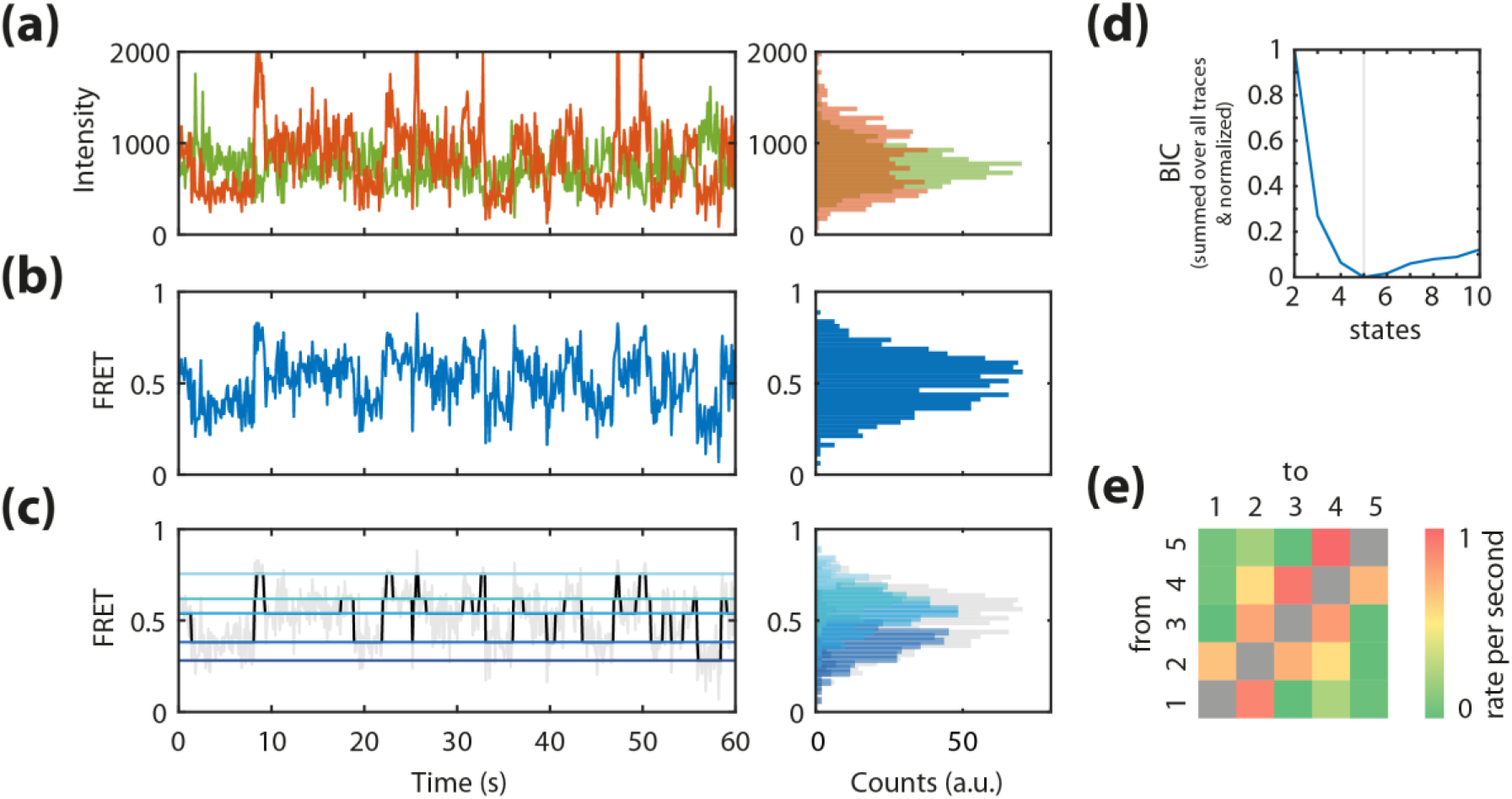
Opening and closing of the *Tm*FtsH domains in the absence of ATP. (a) Representative donor and acceptor fluorescence intensity time trajectories (left panel) and their distributions (right panel). (b) smFRET time trajectory (left panel) constructed from (a) and the derived FRET efficiency histogram (right panel) (c) Viterbi path reconstruction of the smFRET time trajectory in (b) using a five-state model (left panel) and the derived histogram (right panel) (d) Global BIC function. (e) Heatmap of all conformational transition rates of *Tm*FtsH_184,513_ in the absence of ATP.

To resolve the number of states adopted by *Tm*FtsH_184,513_ and to derive kinetic information on inter-domain switching, we analyzed the obtained smFRET time trajectories by global Hidden Markov modelling using ebFRET [34] (Fig. 2c). This analysis revealed that smFRET time traces are best described by five distinct conformational states, as indicated by the minimum in an unbiased global Bayesian Information Criterion (BIC) function (Fig. 2d). A Viterbi path reconstruction of the entire set of time trajectories using a five-state model excellently reproduced the experimental smFRET trajectories (Fig. 2c, left panel and Fig. S7 for more example data) and allowed us to extract state dwell times and kinetic rates. A histogram created from the reconstructed Viterbi paths covered the full distribution of underlying conformational states (Fig. 2c, right panel), substantiating that the conformational switching of *Tm*FtsH_184,513_ is well described by five dynamically interconverting states. A rate matrix (Fig. 2e) generated for all closing and opening transitions revealed that transitions between neighboring states (e.g., state 1 to state 2) are much more frequently observed than transitions between other states (e.g., state 1 to state 4), indicating that conformational switching of *Tm*FtsH_184,513_ occurs primarily in a sequential manner to the nearest neighbor state. A graphical representation summarizing the relative occupancies and kinetic rates for opening and closing transitions between the nearest neighbor states is shown in Figure 3a.

**Figure 3.**
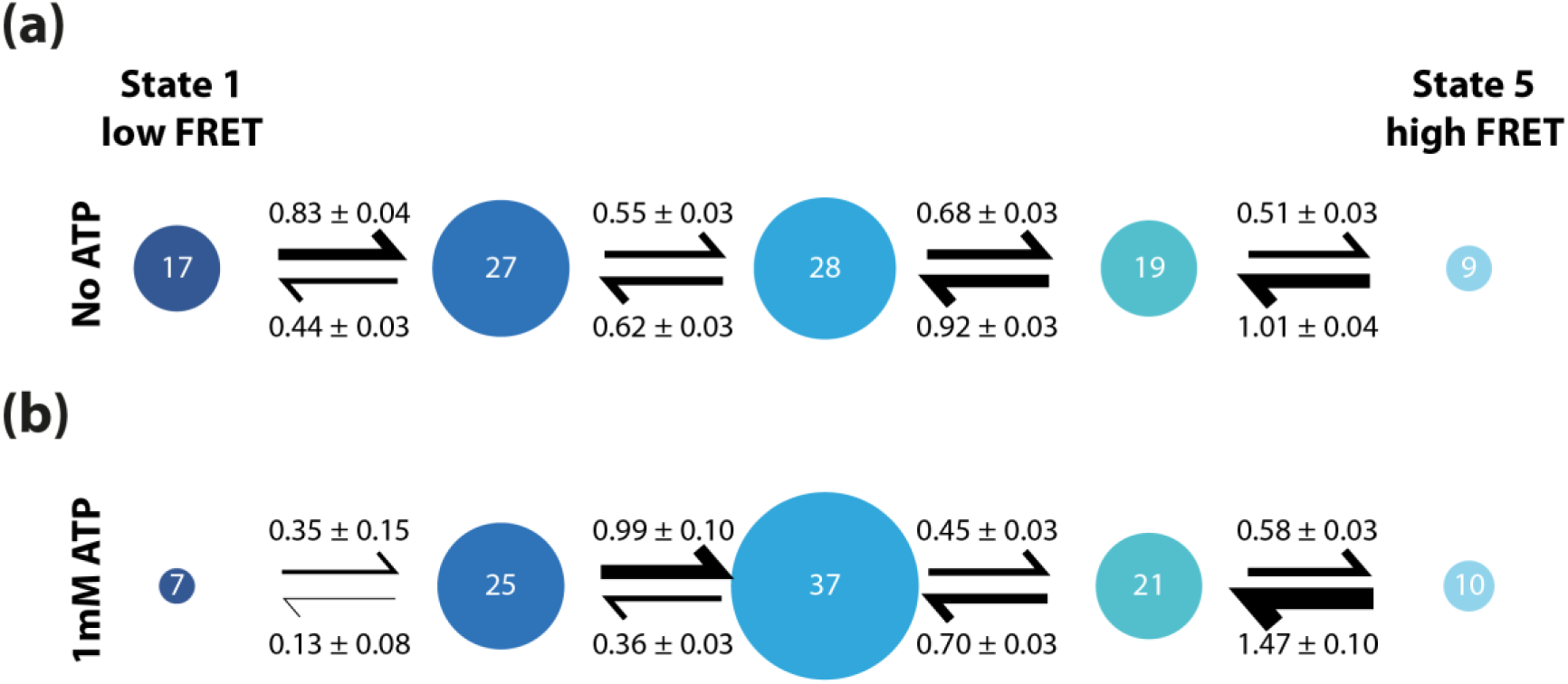
*Tm*FtsH_184,513_ sequential conformational switching without and with ATP. Occupancies (%) and kinetic rates (per second) of *Tm*FtsH_184,513_ without (a) and with 1 mM ATP (b). Size of states and thickness of arrows scale with the occupancy probabilities and kinetic rates, respectively. State occupancies showed an error <3%.

### ATP shifts the population equilibrium towards a more compacted state

In a next step, we wanted to test whether the presence of ATP has an influence on the conformational dynamics of *Tm*FtsH, as ATP has been inferred from crystallographic studies to influence the chemo-mechanical state [19]. To this end, we performed smFRET measurements on *Tm*FtsH_184,513_ in the presence of 1 mM ATP. Similar to our observations without ATP, smFRET time trajectories showed large fluctuations between multiple states on the second timescale (Figs. S4a,b and S7b for more example data). A FRET efficiency histogram created from all trajectories exhibits a broad distribution, indicating again a large opening and closing motion of both domains ranging from *E*_min_ ≈ 0.3 to *E*_max_ ≈ 0.9 (Fig. S4b). Interestingly, the FRET efficiency distribution is skewed to slightly higher FRET efficiencies compared to *Tm*FtsH_184,513_ in the absence of ATP. This indicates that *Tm*FtsH assumes a more compact conformation when ATP is present. To shed light on whether this apparent shift is caused by a repopulation of FRET states or a shift of FRET states towards higher FRET efficiencies, we analyzed the time trajectories using Hidden Markov modelling (*n* = 17 trajectories). As seen for the experiments without ATP, we found conformational transitions between five states indicated by the minimum in the BIC function (Fig. S4d). Moreover, transitions also occurred most frequently between direct neighboring conformational states, which exhibited the highest transition rates (Fig. S4e). The FRET efficiency states in the absence and presence of ATP agree well within error, indicating that similar molecular states are adopted under both conditions (Fig. S4c). However, we observed state population probabilities shifting to a higher population of state 3 as compared to the absence of ATP (Fig. 3b). Thus, while the timescale of interstate conversion is similar, the equilibrium of states is shifted. A higher closing-to-opening ratio between states 1 and 2 and state 2 and 3 in the presence of ATP leads to a depopulation of state 1 and, thus, an increased net probability of the third conformational state. In total, *Tm*FtsH_184,513_ spent 37% of its time in state 3 in the presence of ATP compared to 28% in absence of ATP. Interestingly, the kinetic ratios between state 3 and 4 and state 4 and 5 are similar although absolute numbers vary, thus indicating that ATP does not affect the equilibrium of these transitions to a great extent. Furthermore, we observed that populating state 1 in presence of ATP occurred only in rare cases leading to a 7% occupancy probability (vs. 17% FRET state population without ATP). Taken together, the re-equilibrations explain the overall shift of the open conformational states 1 and 2 towards a more compact conformation in state 3 in the presence of ATP.

### A pathogenic mutation of the human homolog paraplegin hinders *Tm*FtsH compaction

The human homolog of *Tm*FtsH, paraplegin (SPG7), shares 53% structural identity and >70% similarity with *Tm*FtsH [23] (Fig. S3). The pathogenic mutation of paraplegin A510V is associated with the autosomal recessive form of HSP, causing progressive spastic paralysis in the lower limbs [20]. Yet, its molecular effects remain elusive. A510V is highly conserved and located at the hinge interface between the ATPase and protease domains, however, it is far from both active sites [23] and still impairs the function of paraplegin [35]. Hence, we wanted to address if the malfunction caused by the mutation of the *Tm*FtsH-homolog arises from a structural or dynamic deficiency. To this end, we introduced the paraplegin A510V- homologous mutation A359V in *Tm*FtsH_184,513_ and created fluorescently labeled mutant protein, thereafter named *Tm*FtsH_184,513_-A359V. In functional assays, the A359V mutation did not alter ATPase and protease activities compared to *Tm*FtsH_184,513_ (Figs. S1 and S2, respectively). We hexamerized *Tm*FtsH_184,513_-A359V in lipid vesicles and performed smFRET measurements both in the absence and presence of ATP. smFRET time trajectories showed similar transitions between multiple discrete states as observed for *Tm*FtsH_184,513_, revealing that the highly dynamic switching between multiple states on the second timescale is preserved in *Tm*FtsH_184,513_-A359V (Figs. S5a,b, S6a,b). The dynamic behavior resulted also in broad distributions in the cumulative smFRET efficiency histograms ranging between *E*_min_ = 0.2 and *E*_max_ = 0.9 (Figs. S5b, S6b). We analyzed all smFRET trajectories using Hidden Markov modelling, which revealed five conformational states as observed for *Tm*FtsH_184,513_, both in the absence and presence of ATP (Figs. S5c, S6c, S7c,d) and with a preferential switching between neighboring states (Figs. S5d, S6d). In the absence of ATP, the overall state occupancy distribution of *Tm*FtsH_184,513_-A359V is largely unaffected by the point mutation (Fig. 4a) when compared to *Tm*FtsH_184,513_. However, a different behavior between *Tm*FtsH_184,513_-A359V and *Tm*FtsH_184,513_ is observed in the presence of ATP (Fig. 4b). While *Tm*FtsH_184,513_-A359V also undergoes switching on the second timescale, ATP does not induce the overall compaction of *Tm*FtsH towards the conformational state 3 as observed in *Tm*FtsH_184,513_ (c.f. Fig. 3b). Instead, the equilibria between states 1 to 3 were largely unchanged in presence of ATP. This suggests that the conformational switch towards a more closed conformational state in presence of ATP is inhibited in mutant *Tm*FtsH_184,513_-A359V.

**Figure 4.**
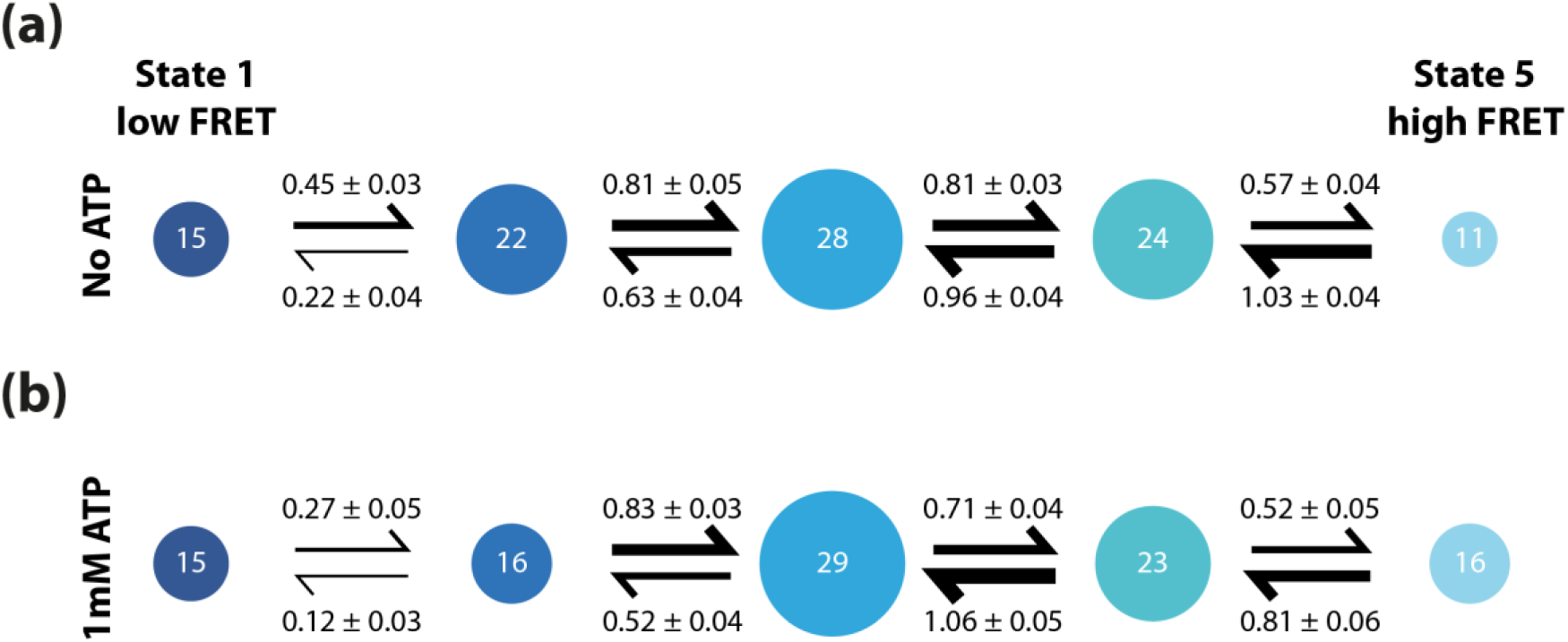
*Tm*FtsH_184,513_-A359V sequential conformational switching without and with 1 mM ATP. Occupancies (%) and kinetic rates (per second) of *Tm*FtsH_184,513_- A359V without (a) and with 1 mM ATP (b). Size of states and thickness of arrows scale with the occupancy probability and kinetic rate, respectively. State occupancies showed an error <3%.

## Discussion

Resolving the inter-domain conformational changes of *Tm*FtsH is an essential step towards elucidating its proteolysis mechanism. Previous crystallographic studies on *Tm*FtsH indicated that binding of ADP drives a large conformational transition from an open state in the apo form to a closed state, in which the ATPase domain is closely associated with the protease domain [10,18,19]. A model of the chemo-mechanical cycle was inferred from these two crystal structures, describing *Tm*FtsH’s function with a power-stroke mechanism, whereby the energy released by ATP hydrolysis is converted into a conformational switch. While these structures granted Ångström-resolved snapshots of two distinct conformers of the protease, they did not reveal the connectivity between the states nor provided time trajectories of structural changes and their coupling to ATP binding.

In this work, we established a kinetic view on *Tm*FtsH’s ATPase and protease inter-domain switch by using time-resolved smFRET to monitor the unsynchronized domain movements of *Tm*FtsH monomers within self-assembled hexamers. We found that *Tm*FtsH is a highly dynamic protease, even in the absence of nucleotide, and undergoes sequential, thermally-driven closing and opening motions through five discrete conformational states with an occupancy on the second timescale. The observed amplitude of the conformational switch between the two domain tips spans a length scale of ∼3–4 nm and is thus consistent with the large-scale hinge-bending motion seen in the crystal structures. Yet, the three additional conformational states, as witnessed by our single-molecule time trace analysis, indicate a much more complex structural reorganization than implied by the two-state picture inferred from the crystallographic snapshots.

In the presence of ATP, *Tm*FtsH also performs highly dynamic switching between five different states on the second timescale but the protease undergoes an inter-domain compaction caused by a depopulation of the open state towards a more closed state. While compaction is in line with the closing action implied by the model derived from the crystal structure upon ADP binding, the thermally driven and, thus, weakly ATP-coupled inter-domain movement behavior markedly differs from the existing conformational model, in which dynamic switching of *Tm*FtsH was described to be purely driven by energy conversion from ATP. Our kinetic data rather suggest a model (Fig. 5a) in which thermal energy, but not chemical energy derived from ATP, provides the major driving force of the conformational fluctuations, while ATP, through a state reequilibration, introduces directionality into these thermal domain motions. Support for a weakly ATP-coupled inter-domain reconfiguration model comes from a comparison of *Tm*FtsH’s ATP consumption rates and its conformational transition rates. The ATPase activity of *Tm*FtsH_184,513_ with 7 × 10^−4^ ATP/monomer/s is significantly slower than the second timescale inter-domain conformational dynamics observed in our smFRET experiments, both without and with ATP, thus substantiating the notion that ATP itself might not be an important driver in conformational changes of *Tm*FtsH in its chemo-mechanical cycle.

**Figure 5.**
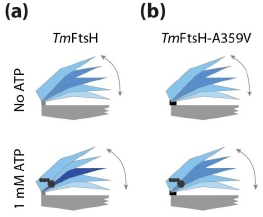
Model of thermally-driven domain motions in *Tm*FtsH and the effect of hindered movement upon introduction of a pathological hinge point mutation. (a) *Tm*FtsH and (b) *Tm*FtsH-A359V without ATP (upper panel) and with 1 mM ATP (bottom panel). Blue scaling of the ATPase domain indicates occupancies of the conformational state, with a darker color indicating a higher occupied state.

A largely energy-independent or weakly ATP-coupled conformational switching mechanism resembles previous work on the dimeric ATPase chaperone Hsp90 [36], where conformational changes between different states of the protein were found to result from thermal fluctuations rather than from the release of chemical energy upon ATP hydrolysis. Low lying energy barriers on the order of a few *k*b*T*s together with multiple discrete conformational states were suggested to drive the conformational changes sequentially in Hsp90. Such a mechanism might also be at play in *Tm*FtsH and explain how thermal fluctuations are sufficient for *Tm*FtsH to drive sequential transitions from one state to a neighbor state, thus not necessitating ATP energy conversion for its domain fluctuations.

From a functional point of view, the mechanism of thermally-induced conformational switching in *Tm*FtsH may be of importance for its unfoldase or translocase activity. By severing the large conformational switching into small transitions, the partial thermally-unfolded protein substrate bound at the ATPase domain of *Tm*FtsH could be transferred to the proteolytic domain, similar to previous reports of thermal ratchets [37,38], while ATP might aid in establishing directionality in this process. Even though conformational switching appears largely independent of energy use, we speculate that ATP hydrolysis might be required for unfolding of mechanically stable substrates, as recently observed for the bacterial ClpXP system and I27 as well as green fluorescent protein (GFP) unfolding [6,8,39–41].

The thermally-driven domain motions in *Tm*FtsH suggest an important role of the hinge interface connecting the ATPase and protease domain in mediating the conformational switch, which might be disturbed when mutations are introduced. To address this question, we studied the conformational dynamics of *Tm*FtsH carrying the A359V mutation, a hinge point mutation which is homologous to the HSP-pathogenic mutation of paraplegin A510V [20,23]. We found that *Tm*FtsH-A359V also performs switching between five different conformational states with thermally driven interconversion on the seconds timescale. However, *Tm*FtsH-A359V did not show the characteristic compaction towards state 3 upon addition of ATP as seen in *Tm*FtsH, but exhibited, within error, unchanged population occupancies of its five states. With 4 × 10^−4^ ATP/hexamer/s, the point mutation itself did not significantly alter the ATPase activity of *Tm*FtsH_184,513_-A359V. Thus, we speculate that the mutation A359V might hinder an allosteric communication upon ATP binding or hydrolysis as the molecule is ‘locked’ in a more open state than could be evoked by ATP binding or hydrolysis. We anticipate that this hindered communication could also be the molecular cause of the malfunction in the human paraplegin A510V mutant (Fig. 5b).

Whereas the presence of substrate stimulates ATP hydrolysis (Fig. S1), it remains unknown if substrate binding would change the occupancies of the molecular conformations or the dynamic behavior of domain motions and their ATP coupling behavior. Further smFRET experiments using different protease substrates with both *Tm*FtsH and *Tm*FtsH-A359V should shed light on the existence of a substrate-induced energy-coupled mechanism of degradation and potentially further malfunctions of pathogenic mutations. Noteworthy, our results, together with earlier work that reported ATP-independent proteolytic activity of *Tm*FtsH—which we also observed with the weakly folded protein substrate casein for *Tm*FtsH_184,513_ (Fig. S2)—indicate the possibility that the proteolytic activity could be decoupled from ATPase activity. Thus, it remains to be explored how *Tm*FtsH could use a dual mechanism for processing its protein substrates: one ATP dependent and one ATP independent.

In conclusion, we found that conformational changes between the ATPase and the protease domain of *Tm*FtsH are driven by thermal motions and only weakly coupled to ATP. *Tm*FtsH adopts five discrete, well-defined states during closing and opening cycles, which remain to be structurally resolved. The presence of ATP favors compaction of *Tm*FtsH with higher occupancy of state 3 along the closing cycle, yet all five conformational states are frequently adopted by *Tm*FtsH. Introducing a mutation in the hinge interface of *Tm*FtsH, homologous to a human pathogenic mutation of paraplegin causing HSP, prevents the compaction observed by the ATPase and protease domains of *Tm*FtsH, a perturbation that might explain the pathogenic activity of the human homolog paraplegin causing HSP.

## Materials and Methods

### Protein design, production, and purification

The cDNA sequence corresponding to amino acid residues 147–610 of *Tm*FtsH was inserted between the NcoI and NotI sites of a pET28a(+) vector (Novagen) to encode an N-terminal His_6_-tagged *Tm*FtsH fusion protein lacking the transmembrane domain (Δtm) of *Tm*FtsH (His_6_-(Δtm)*Tm*FtsH(147–610)), as described in Bieniossek et al. [18,19]. A plasmid encoding a double Cys variant (*Tm*FtsH_184,513_) of His_6_- (Δtm)*Tm*FtsH(147–610) for site-specific labeling with thiol-reactive fluorophores at positions 184 and 513 was created by introducing a non-native Cys at position 184 and by replacing the two native Cys residues at positions 255 and 564 by Ser residues to prevent unspecific labeling. The expression vector encoding the mutant *Tm*FtsH_184,513_-A359V variant was generated from the *Tm*FtsH_184,513_ plasmid by an Ala-to-Val replacement at position 359. Mutations were introduced using the QuikChange Lightning Multi site-directed mutagenesis kit (Novagen). Target mutations in all constructs were confirmed by DNA sequencing.

Recombinant production of *Tm*FtsH_184,513_ and *Tm*FtsH_184,513_-A359V fusion proteins was performed using the *Escherichia coli* host strain BL21(DE3) (Novagen). Briefly, bacterial cells were transformed with the respective *Tm*FtsH_184,513_- or *Tm*FtsH_184,513_-A359V-encoding pET28a(+) expression vector and grown in Luria-Bertani medium (supplemented with 50 µg/ml Kanamycin) to an optical density at 600 nm (OD_600_) of ∼0.6. Expression was induced by the addition of isopropyl-*β*-D-1-thiogalactopyranoside (IPTG) to a final concentration of 0.5 mM and cells cultivated for a further 3 h at 37°C. Cells were harvested by centrifugation and lysed using an Emulsiflex high-pressure homogenizer (Avestin). The soluble crude extract was then separated from the cell debris and insoluble content by centrifugation for 40 min at 40,000 rpm at 4°C. The supernatant was subjected to heat purification at 75°C for 3 min, followed by subsequent centrifugation for 10 min at 40,000 rpm and 4°C, according to previously published procedures [18,19].

Protein purification was performed via immobilized-metal-ion affinity chromatography (IMAC). Briefly, the soluble fraction of the cell lysate was applied to a HisTrap FastFlow column (GE Healthcare) that had been pre-equilibrated with binding buffer (20 mM HEPES pH 8.0, 300 mM NaCl, 10 mM imidazole). After extensive washing with washing buffer (20 mM HEPES pH 7.5, 300 mM NaCl, 40 mM imidazole), his-tagged proteins were eluted with elution buffer (20 mM HEPES pH 7.5, 300 mM NaCl, 250 mM imidazole).

### ATPase and protease activity assays

ATPase activities of *Tm*FtsH_184,513_ and *Tm*FtsH_184,513_- A359V were tested using the EnzChek Phosphate Assay kit (Thermo Fisher) according to the manufacturer’s instructions. Absorption was monitored at 360 nm (Fig. S1) after successively adding ATP (1 mM), cI–ssrA substrate (0.8 μ M) and *Tm*FtsH_184,513_ (14.4 μ M) or *Tm*FtsH_184,513_- A359V (14.4 μ M) to the blank buffer solution containing 20 mM HEPES pH 8.0, 150 mM KCl, 10% glycerol, 5 mM MgOAc, 12.5 µM ZnOAc. The cI–ssrA substrate, an N-terminal His_6_- tagged and C-terminal ssrA-tagged (AANDENYALAA) repressor protein cI fusion protein, was recombinantly produced in *E. coli* from a pET28(+) vector and purified by IMAC, followed by dialysis against buffer (20 mM HEPES pH 8.0, 150 mM KCl, 10% glycerol).

Protease activity tests of *Tm*FtsH_184,513_ and *Tm*FtsH_184,513_-A359V were carried out by incubating 2.4 µM protease sample with various combinations of 1 mM casein, 1 mM ATP and 1 mM EDTA overnight at 24°C and 50°C in buffer containing 20 mM HEPES pH 8.0, 150 mM KCl, 10% glycerol, 5 mM MgOAc, 12.5 µM ZnOAc. Protease activity was monitored by substrate degradation using sodium dodecyl sulfate polyacrylamide gel electrophoresis (SDS-PAGE) (Fig. S2). Casein stock solution (0.65% (*w*/*v*)) was prepared by dissolving bovine casein (Sigma) in buffer containing 50 mM HEPES pH 8.0.

### Protein labeling

*Tm*FtsH_184,513_ and *Tm*FtsH_184,513_-A359V were labeled with maleimide-functionalized sulfo-Cy3 and sulfo-Cy5 dyes (both from GE Healthcare). Purified *Tm*FtsH_184,513_ or *Tm*FtsH_184,513_-A359V at a final concentration of 20 µM were reacted with a 10–15-fold excess of both dyes under reducing conditions (0.4 mM tris(2-carboxyethyl)phosphine (TCEP)) in buffer containing 20 mM HEPES pH 7.5, 300 mM NaCl, 250 mM imidazole. The labeling reaction was carried out for 2 h at room temperature, followed by overnight incubation at 4°C. Labeled protein was separated from the unreacted dyes by IMAC using a spin-column purification protocol. Briefly, after diluting the labeling reaction tenfold in binding buffer (20 mM HEPES pH 8.0, 300 mM NaCl, 10 mM imidazole), the solution was applied to a Ni-NTA Agarose resin (Thermo Fisher) that had been pre-equilibrated with binding buffer. After, extensive spin-washing at 2,000 rpm with binding buffer until the supernatant was clear and free of residual dyes, the labeled protein was eluted with elution buffer (20 mM HEPES pH 8.0, 300 mM NaCl, 250 mM imidazole) by centrifugation for 2 min at 2,000 rpm. Labeling efficiency was determined photometrically.

### Sample preparation for single-molecule FRET experiments

DMPC and biotin-DPPE lipid powders (both from Avanti Polar Lipids) were dissolved in chloroform and mixed in a 99:1 DMPC:biotin-DPPE ratio at a final concentration of 5 mg/ml. Solvent was subsequently removed by vacuum drying overnight to yield a dried lipid cake. In a separate step, labeled *Tm*FtsH_184,513_ was mixed with unlabeled *Tm*FtsH_184,513_ in a 1:5 ratio at a final protein concentration of 12 µM in buffer (20 mM HEPES pH 8.0, 150 mM KCl, 10% glycerol, 5 mM MgOAc, 12.5 µM ZnOAc) using labeled and unlabeled protein from the same purification batch. The protein mix was then added to the dried lipid cake and incubated at 37°C for 30 min. After addition of additional buffer (20 mM HEPES pH 8.0, 150 mM KCl, 10% glycerol, 5 mM MgOAc, 12.5 µM ZnOAc), protein encapsulation into unilamellar lipid vesicles was performed by 35-fold extrusion of the protein–lipid suspension through polycarbonate filters with a pore diameter of 200 nm using a Mini-Extruder (Avanti Polar Lipids) at 37°C according to the manufacturer’s instructions. *Tm*FtsH_184,513_-loaded vesicles were then immobilized on a biotinylated PEG-coated flow chamber via a biotin–neutravidin–biotin sandwich. Before single-molecule imaging, the sample chamber was supplemented with imaging buffer (20 mM HEPES pH 8.0, 100 mM NaCl, 5 mM MgOAc, 12.5 μ M ZnOAc) containing saturated, aged Trolox (6-hydroxy-2,5,7,8-tetramethylchroman-2-carboxylic acid, Sigma) and an oxygen scavenging system (100 μg/ml glucose oxidase, 0.8% (*w*/*v*) D-glucose, and 1 μU/ml catalase). For measurements in the presence of ATP, 1 mM ATP was added to the imaging buffer solution. Because the transition temperature of DMPC is at room temperature (∼24°C)—the temperature at which our experiments were performed—coexistence of liquid and gel phases of the lipids permeabilizes the membrane for exchanging small molecules such as ATP [30]. Sample preparation for experiments with *Tm*FtsH_184,513_-A359V was carried out in the same way.

### Single-molecule FRET experiments

A custom-built prism-type TIRF microscope was used for single-molecule data acquisition as previously described [31,32,42]. Imaging was performed at room temperature. Single-molecules donor and acceptor intensities (*I*_*D*_, *I*_*A*_) were recorded over time at 100-ms time resolution and smFRET time trajectories were extracted by calculating *E*app for each collected data point (see Results). Distances (*r*) were approximated on the basis of the Förster equation, *r* = *R*0 × (1/(*E* – 1))^1/6^, where *E* is the corrected FRET efficiency, and *R*0 is the Förster distance of the Cy3/Cy5 FRET pair (*R*0 = 5 nm), with the assumption that the fluorophores can freely rotate at the labeling site. Only traces that exhibited single donor and acceptor bleaching steps were evaluated. Hidden-Markov model analysis was performed using ebFRET as described [34] (https://ebfret.github.io).

## Acknowledgements

We thank all members of the Schlierf lab for discussions. In particular, the authors would like to thank Andreas Hartmann for help in data acquisition and establishing data analysis software. This study was supported by the German Ministry for Science and Education BMBF (grant 03Z2EN11 to MS), the Deutsche Forschungsgemeinschaft (grants SCHL1896/2-1 and SCHL1896/3-1 to MS) and Stipendienstiftung Rheinland-Pfalz with a scholarship to GK.

## Supplementary data

Supplementary data accompanying this manuscript: ATPase activity assay of *Tm*FtsH_184,513_ and *Tm*FtsH_184,513_-A359V; Protease activity assay of *Tm*FtsH_184,513_ and *Tm*FtsH_184,513_-A359V; structural alignment of *Tm*FtsH and its structural human homolog paraplegin; smFRET data on *Tm*FtsH_184,513_ with 1 mM ATP, *Tm*FtsH_184,513_-A359V without ATP, *Tm*FtsH_184,513_-A359V with 1 mM ATP; selection of smFRET time trajectories of *Tm*FtsH_184,513_ and *Tm*FtsH_184,513_-A359V in the presence and absence of ATP.

## Supplementary Data

### Supplementary Figures

**Figure S1.**
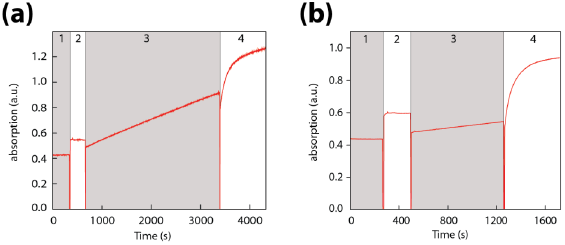
ATPase activity assay of *Tm*FtsH_184,513_ (a) and *Tm*FtsH_184,513_-A359V (b). Monitored is the absorbance of 2-amino-6-mercapto-7-methyl-purine at 360 nm that is generated in the EnzChek Phosphate Assay upon release of free phosphate in solution. The four phases indicate (1) the background absorption of the buffer (20 mM HEPES pH 8.0, 150 mM KCl, 10% glycerol, 5 mM MgOAc, 12.5 µM ZnOAc), (2) the absorbance upon addition of 1 mM ATP, (3) the absorbance upon addition of 14.4 µM *Tm*FtsH_184,513_ or *Tm*FtsH_184,513_-A359V, and (4) the absorbance upon addition of 0.8 µM cI–ssrA substrate. The reactions were performed at room temperature.

**Figure S2.**
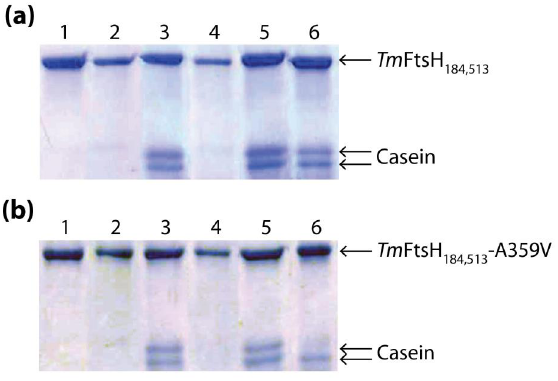
Protease activity assay of *Tm*FtsH_184,513_ and *Tm*FtsH_184,513_-A359V. (a) *Tm*FtsH_184,513_ (2.4 µM) incubated with 1 mM casein and 1 mM ATP at 24°C (1) and at 50°C (2); with 1 mM casein only at 24°C (3) and at 50°C (4); and with 1 mM casein, 1 mM ATP, and 0.1 mM EDTA at 24°C (5) and at 50°C (6). (b) *Tm*FstH_184,513_-A359V (2.4 µM) incubated with 1 mM casein and 1 mM ATP at 24°C (1) and at 50°C (2); with 1 mM casein only at 24°C (3) and at 50°C (4) and with 1 mM casein, 1 mM ATP, and 0.1 mM EDTA at 24°C (5) and at 50°C (6). The reaction was conducted overnight in buffer containing 20 mM HEPES pH 8.0, 150 mM KCl, 10% glycerol, 5 mM MgOAc, 12.5 µM ZnOAc.

**Figure S3.**
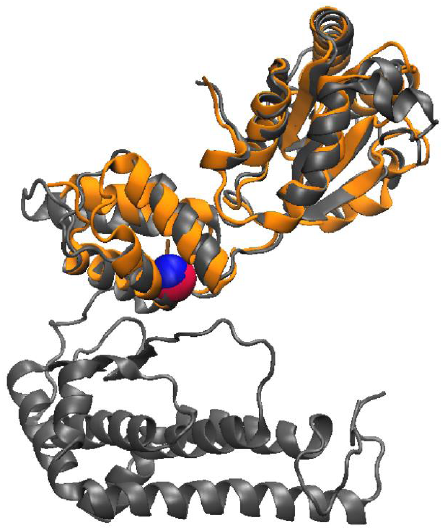
Structural alignment of *Tm*FtsH (gray; pdb: 3KDS) and its structural human homolog paraplegin (orange; pdb: 2QZ4). Alanine residue 359 (blue, C_α_) and the homologous pathogenic mutation A510V (red, C_α_) are highlighted. Both mutations are located at the boundary of the ATPase and protease domains. Alignment was performed in VMD.

**Figure S4.**
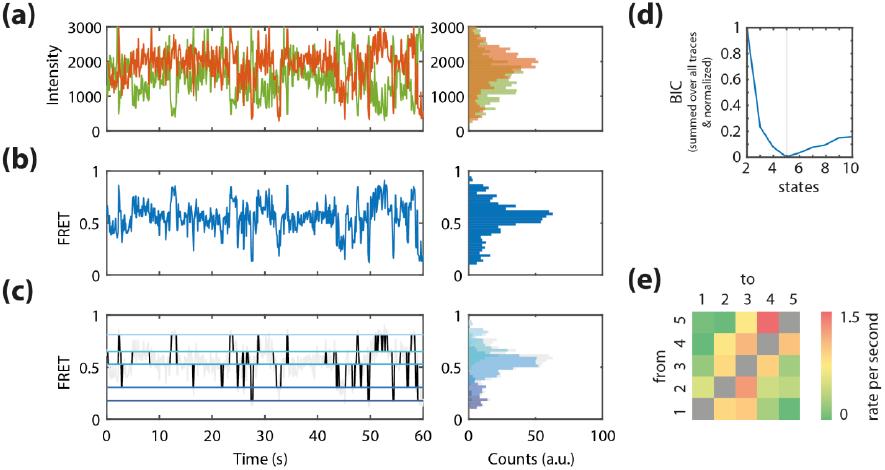
*Tm*FtsH_184,513_ with 1 mM ATP. (a) Representative donor and acceptor fluorescence intensity time trajectories (left panel) and their distributions (right panel). (b) smFRET time trajectory (left panel) constructed from (a) and the derived FRET efficiency histogram (right panel). (c) Viterbi path reconstruction of the smFRET time trajectory in (b) using a five-state model (left panel) and the derived histogram (right panel). (d) Global BIC function. (e) Heatmap of all conformational transition rates of *Tm*FtsH_184,513_ in the presence of ATP.

**Figure S5.**
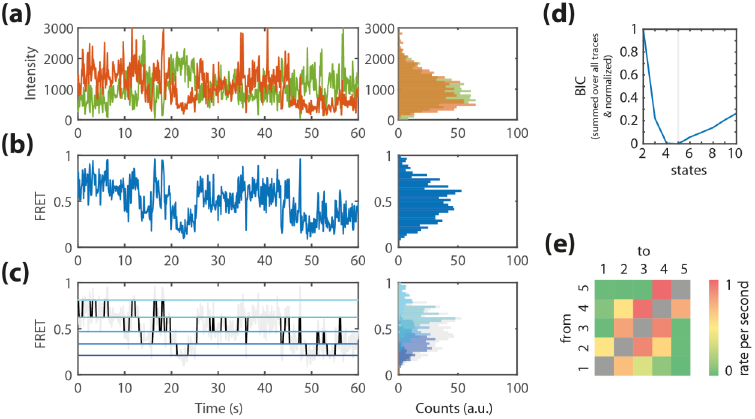
*Tm*FtsH_184,513_-A359V without ATP. (a) Representative donor and acceptor fluorescence intensity time trajectories (left panel) and their distributions (right panel). (b) smFRET time trajectory (left panel) constructed from (a) and the derived FRET efficiency histogram (right panel). (c) Viterbi path reconstruction of the smFRET time trajectory in (b) using a five-state model (left panel) and the derived histogram (right panel). (d) Global BIC function. (e) Heatmap of all conformational transition rates of *Tm*FtsH_184,513_-A359V in the absence of ATP.

**Figure S6.**
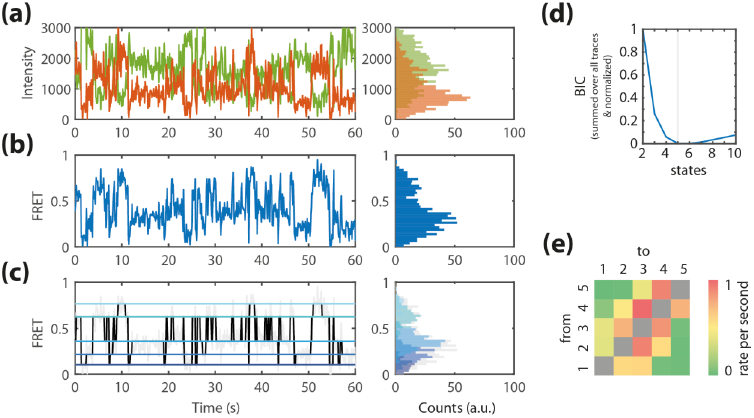
*Tm*FtsH_184,513_-A359V with 1 mM ATP. (a) Representative donor and acceptor fluorescence intensity time trajectories (left panel) and their distributions (right panel). (b) smFRET time trajectory (left panel) constructed from (a) and the derived FRET efficiency histogram (right panel). (c) Viterbi path reconstruction of the smFRET time trajectory in (b) using a five-state model (left panel) and the derived histogram (right panel). (d) Global BIC function. (e) Heatmap of all conformational transition rates of *Tm*FtsH_184,513_-A359V in the presence of ATP.

**Figure S7.**
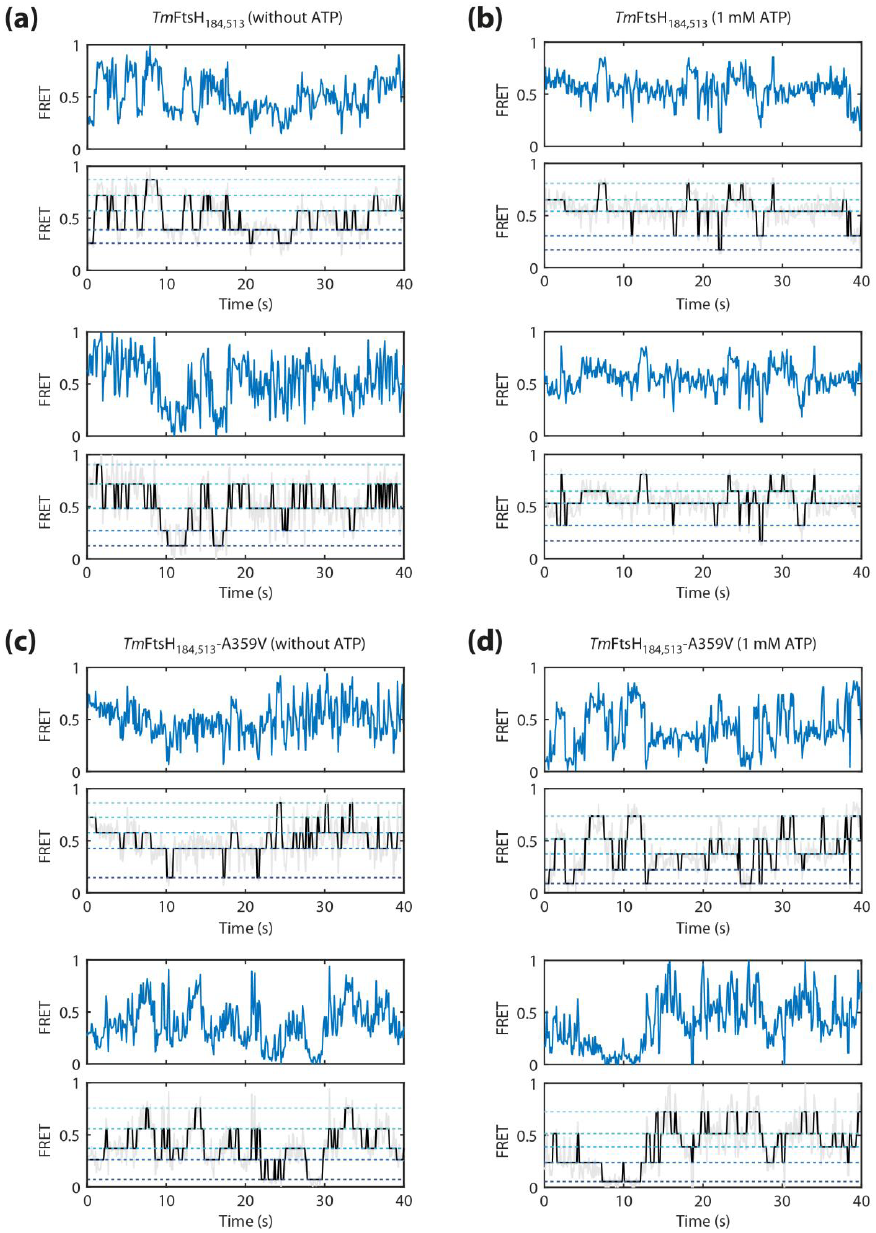
Selection of smFRET time trajectories of *Tm*FtsH_184,513_ and *Tm*FtsH_184,513_-A359V in the presence and absence of ATP. Top panels: smFRET time trajectories. Bottom panels: Viterbi path reconstruction of the smFRET trajectories using a five-state model.

## Abbreviations

AAA+: ATPases associated with diverse cellular activities
*Tm*FtsH: *Thermotoga maritima* FtsH
HSP: hereditary spastic paraplegia
smFRET: single-molecule Förster resonance energy transfer
BIC: Bayesian Information Criterion

